# Various miRNAs are involved in efficient HCV replication

**DOI:** 10.1101/2020.01.10.901488

**Authors:** Chikako Ono, Takasuke Fukuhara, Songling Li, Jian Wang, Asuka Sato, Takuma Izumi, Yuzy Fauzyah, Takuya Yamamoto, Yuhei Morioka, Nikolay V. Dokholyan, Daron M. Standley, Yoshiharu Matsuura

## Abstract

One of the determinants for tissue tropism of hepatitis C virus (HCV) is miR-122, a liver-specific microRNA. Recently, it has been reported that interaction of miR-122 to HCV RNA induces a conformational change of the 5’UTR internal ribosome entry site (IRES) structure to form stem-loop II structure (SLII) and hijack of translating 80S ribosome through the binding of SLIII to 40S subunit, which leads to efficient translation. On the other hand, low levels of HCV-RNA replication have also been detected in some non-hepatic cells; however, the details of extrahepatic replication remain unknown. These observations suggest the possibility that miRNAs other than miR-122 can support efficient replication of HCV-RNA in non-hepatic cells. Here, we identified a number of such miRNAs and show that they could be divided into two groups: those that bind HCV-RNA at two locations (miR-122 binding sites I and II), in a manner similar to miR-122 (miR-122-like), and those that target a single site that bridges sites I and II and masking both G28 and C29 in the 5’UTR (non-miR-122-like). Although the enhancing activity of these non-hepatic miRNAs were lower than those of miR-122, substantial expression was detected in various normal tissues. Furthermore, structural modeling indicated that both miR-122-like and non-miR-122-like miRNAs not only can facilitate the formation of an HCV IRES SLII but also can stabilize IRES 3D structure in order to facilitate binding of SLIII to the ribosome. Together, these results suggest that HCV facilitates miR-122-independent replication in non-hepatic cells through recruitment of miRNAs other than miR-122. And our findings can provide a more detailed mechanism of miR-122-dependent enhancement of HCV-RNA translation by focusing on IRES tertiary structure.

**Author summary:** One of the determinants for tissue tropism of hepatitis C virus (HCV) is miR-122, a liver-specific microRNA, which is required for efficient propagation. Recently, it has been reported that interaction of miR-122 with the 5’UTR of HCV contributes to the folding of a functional IRES structure that is required for efficient translation of viral RNA. In this study, we examined the minimum motifs in the seed region of miRNAs required for the enhancement of HCV replication. As a result, we found two groups of non-hepatic miRNAs: “miR-122-like miRNAs” that can bind HCV-RNA at two locations in a manner similar to miR-122, and “non-miR-122-like miRNAs” that target a single site that masking both G28 and C29 in the 5’UTR. The interaction of these non-hepatic miRNAs with the 5’UTR can facilitate not only the folding of active HCV IRES but also the stabilization of IRES 3D structure in order to facilitate binding to the ribosome. These results suggest the possibility of replication of HCV in non-hepatic cells through interaction with miRNAs other than miR-122 and provide insight into the establishment of persistent infection of HCV in non-hepatic tissues that lead to the development of extrahepatic manifestations.

## Introduction

Hepatitis C virus (HCV) infects over 71 million people worldwide and is a major cause of chronic hepatitis, liver cirrhosis and hepatocellular carcinoma [1]. One of the most important host factors for HCV infection is a liver-specific microRNA (miRNA), miR-122 [2]. On the other hand, chronic infection with HCV is often associated with extrahepatic manifestations such as mixed cryoglobulinemia, B-cell lymphoma, thyroiditis, and diabetes mellitus [3]. Supported by clinical observations, low levels of replication of HCV-RNA were detected in PBMCs and neuronal tissues in chronic hepatitis C patients [4, 5] and the treatment of chronic hepatitis C patients who developed B cell lymphoma by direct-acting antivirals for HCV resulted in the clearance of HCV and lymphoma [6]. These observations suggest that the replication of HCV in non-hepatic cells can be established in miR-122-deficient condition.

In general, miRNAs negatively regulate translation of target mRNA through interaction with the 3’UTR in a sequence-specific manner. In this way, miR-122 regulates the expression of genes involved in the maintenance of liver homeostasis, including lipid metabolism, iron metabolism, and carcinogenesis [7, 8]. In contrast, miR-122 has also been shown to stabilize HCV-RNA [9] and enhance internal ribosome entry site (IRES)-mediated translation [2, 10, 11] and replication [12] of HCV-RNA through direct interaction with the 5’UTR of HCV [13, 14]. The HCV 5’UTR has two binding sites (sites I and II), which are highly conserved among HCV genotypes [15], to which the miR-122 seed sequence can bind [2, 13]. In addition, the overhanging regions in miR-122 have been reported to interact with HCV-RNA and are important for HCV-RNA abundance [14]. Importantly, activation of translation via IRES is promoted by the Argonaut-containing miRNA-induced silencing complex (miRISC) [11]. The miR-122-miRISC complex prevents degradation of HCV-RNA by the cellular 5’-3’exonucleases Xrn1 and Xrn2, and stabilizes the HCV-RNA interaction [16-18]. Upon HCV infection, the interaction between miR-122-miRISC and HCV-RNA further results in the sequestration of miR-122 from host mRNA targets, a phenomenon known as the “sponge effect”, which may be responsible for the long-term oncogenic potential of HCV infection [19]. Recently, it has been reported that miR-122 has an RNA chaperone-like function that induces the folding of the HCV IRES such that in it can readily associate with the 80S ribosome for efficient translation [20, 21]. The functional HCV IRES consists of three parts: the long arm, the short arm and the body. In the liver, HCV hijacks the translating 80S ribosome by using miR-122 to alter the fold of the IRES body in order to bind to the 40S platform [22]. However, it is currently unknown how persistent infection in non-hepatic cells, which are deficient in miR-122, is established.

Previously, we and other groups have reported that an adaptive mutant possessing a G28A substitution in the 5’UTR of the HCV genotype 2a emerges during serial passages of wild type HCV in miR-122 knockout cells, and that these mutants exhibit efficient RNA replication in the absence of miR-122 [23, 24]. Recently, it has been reported that the G28A mutation in the 5’UTR alters the energetics of IRES folding in manner that is miR-122-independent but structurally similar to that of miR-122-mediated folding [20, 21]. Interestingly, the G28A mutation does not occur under miR-122-abundant conditions. Moreover, in clinical samples obtained from patients infected with HCV genotype 2, the rate of G28A mutation was higher in miR-122-deficient peripheral blood mononuclear cells (PBMCs) than in sera that mainly included hepatocyte-derived HCV [23]. Therefore, in this study, we examined the possibility of HCV-RNA replication in miR-122-deficient cells, such as non-hepatic cells, and of participation of miRNAs other than miR-122 to support replication of HCV-RNA.

By mutagenesis analysis, we observed miRNA binding in a manner distinct from that of miR-122 that resulted in enhanced HCV-RNA replication. While miRNA-122 binds HCV-RNA at sites I and II using 6 matching nucleotides containing a GAGUG motif in the seed sequence, we observed a pattern wherein one miRNA molecule was bound to a position intermediate to sites I and II using 7 nucleotides. Although the ability of such miRNAs to promote HCV-RNA replication in miR-122 knockout cells was lower than that of miR-122, the expression of the miRNAs significantly enhanced viral RNA replication. Furthermore, substantial expression of the miRNAs was observed in various human tissues; expression of the miRNAs enhanced HCV propagation in miR-122 knockout cells and increased viral titers by serial passages without emergence of the G28A mutation. These results suggest that HCV facilitates persistent infection in miR-122-deficient non-hepatic cells through emergence of mutant viruses and recruitment of miRNAs other than miR-122, which leads to development of extrahepatic manifestations in chronic hepatitis C patients.

## Results

### Interactions between miR-122 and 5’UTR required for enhancement of HCV-RNA replication

The direct interaction of two molecules of miR-122 with the 5’UTR is known to be essential for efficient HCV-RNA replication in hepatocytes (Fig 1A). One miR-122 molecule binds the HCV-RNA at positions 1-3 and 21-27 using miRNA positions 15-17 and 2-8, respectively, where the latter position, is called site I. A second miR-122 molecule binds the HCV-RNA at positions 29-32 and 37-42 using miRNA positions 13-16 and 2-7, respectively, where the latter position is called site II. To quantify the contribution of specific nucleotides in miR-122 to the enhancement of HCV-RNA replication, a library of miR-122 point-mutants was employed (Fig 1B). Variants of miR-122 possessing substitutions in the seed region and a plasmid encoding an infectious cDNA of HCV, pHH-JFH1mt, were transfected into miR-122 deficient (751-122KO) cells and intracellular HCV-RNA was determined at 3 days post-transfection. Expression of variants containing mutations at positions 1, 8 and 16 exhibited efficient enhancement of HCV-RNA replication, comparable to that of wild type miR-122 (930-fold increase) (Fig 1C). In contrast, mutation of positions 3 to 7 severely impaired HCV-RNA replication. Mutation of positions 2 and 15 resulted in 3-fold and 13.5-fold increase in HCV-RNA replication relative to control miRNA, respectively. We also examined whether the interaction of miR-122 at nucleotide positions 3-7 in the seed region (GAGUG motif, red characters in Fig 1A) was sufficient to enhance HCV-RNA replication or not. The expression of a miR-122 mutant possessing the GAGUG motif with only a 5-nucleotide match, miR-122-GAGUG, couldn’t enhance HCV-RNA replication significantly in 751-122KO cells (S1 Fig), suggesting that an miRNA possessing the GAGUG motif with a 6-nucleotide match in the seed region, such as where positions 2 or 8 were mutated, would be capable of enhancing of HCV-RNA replication.

**Fig 1.**
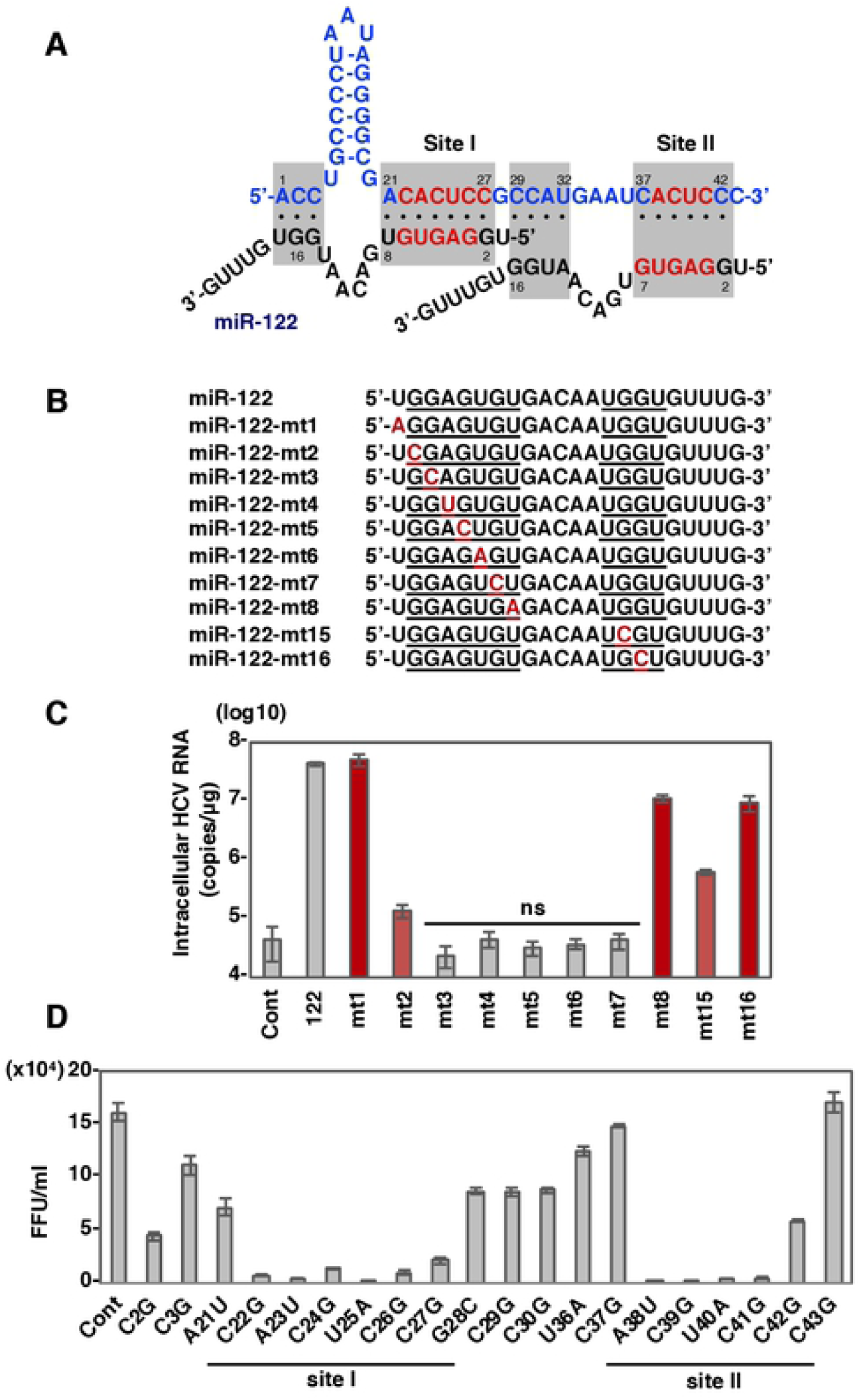
Identification of core sequence of miR-122 and 5’UTR of HCV-RNA required for the enhancement of viral RNA replication. (A) Diagrams of possible interaction between HCV 5’UTR and miR-122. (B) Sequence alignment of miR-122 and its mutant derivatives, miR-122-mt1, -mt2, -mt3, -mt4, -mt5, -mt6, -mt7, -mt8, -mt15 or -mt16. (C) Intracellular HCV-RNA levels of 751-122KO cells infected with JFH1 in the presence of mimic control, miR-122 or its mutant derivatives were determined at 72 hpi by qRT-PCR. (D) Various pHH-JFH1 plasmids encoding HCV mutants at miR-122 interaction sites were transfected into Huh7.5.1 cells and infectious titers in the culture supernatants at 3 dpi were determined by focus formation assay. Error bars indicate the standard deviation of the mean and asterisks indicate significant differences (*P < 0.05; **P < 0.01) versus the results for the control.

To further examine the effects of the mutations on HCV propagation, the mutant plasmids of pHH-JFH1mt were transfected into Huh7.5.1 cells and infectious titers in the culture supernatants were determined at 4 days post-transfection. HCV mutants possessing substitutions C22G, A23U, C24G, U25A, C26G, and C27G in site I, and A38U, C39G, U40A, and C41G in site II (nucleotides of HCV RNA shown in red in Fig 1A) significantly reduced infectious titers to less than 20% of wild type (Fig 1D). Interestingly, HCV mutants C2G, C3G, A21U, G28C, C29G, C30G, U36A, C37G, C42G, and C43G exhibited substantial replication, consistent with the rescue of replication of HCV-RNA in 751-122KO cells by the expression of miR-122 variants containing mutations at positions 1, 8, 15, or 16 (Fig 1C). Nucleotides G16 and G15 in miR-122 have been shown to be required for enhanced replication of HCV-RNA by 3’ overhanging to nucleotides C2 and C3 of HCV RNA, respectively [14]. In addition, 5’RACE analysis revealed that a revertant virus possessing a G to C substitution has emerged after 2 rounds of passage of the C3G mutant (S2 Fig). We consistently observed a factor of 10 reduction in HCV-RNA replication upon mutation G15C in miR-122 (Fig 1C), suggesting that a cytosine at position of 3 in the HCV RNA may be required for efficient propagation of HCV through interaction with miR-122. Collectively, these results suggest that nucleotide positions 22-27 in site I and 38 to 41 in site II play important roles in efficient replication of HCV through interaction with miR-122.

### Identification of miR-122-like miRNAs that enhance HCV-RNA replication

To find miRNAs other than miR-122 that could facilitate enhancement of HCV-RNA replication, we searched the miRBase database (http://www.mirbase.org), which contained 2656 human miRNAs, for miRNAs able to target positions 22-27 and 38-41 of HCV-RNA with a 6-nucleotide match that includes a GAGUG motif in the seed region. We identified a single miRNA (miR-504-3p) that matched all the search criteria (S3A Fig). We also identified 6 additional miRNAs (miR-574-5p, miR-3659, miR-1236-5p, miR-4481, miR-4745-5p and miR-4765) that possessed 5 complementary nucleotides in both site I and II but matched the other search criteria (S3A Fig). Among these miRNAs, we focused on 3 candidates: miR-504-3p, due to its high similarity to miR-122; miR-574-5p, because it has been reported to play a functional role in neuronal cells [33, 34]; miR-1236-5p, because it has substantial expression in several tissues [34]. The tissue specificity index (TSI) can be used classify miRNAs as tissue-specific (TSI>0.85) or housekeeping (TSI<0.5) based on the expression pattern [34]. The TSI of miR-122 is 0.97, while those of miR-574-5p and miR-1236-5p are 0.79 and 0.68, respectively (S3A Fig). To confirm that the candidate miRNAs suppress translation of endogenous mRNA, a pmirGLO vector carrying the complementary sequence of each miRNA under the luciferase coding sequence was transfected into 751-122KO cells together with the corresponding miRNA mimics. Luciferase activities in 751-122KO cells transfected with the miRNA mimics of miR-122, miR-574-5p, and miR-1236-5p were strongly suppressed, while those of miR-504-3p were weakly but significantly reduced compared with those of control mimics (S3B Fig).

Next, to examine the effect of the candidate miRNAs on the enhancement of HCV-RNA replication, 751-122KO cells transfected with each miRNA mimic were infected with HCV 12 hours post-transfection and intracellular HCV-RNA was determined at 72 hours post-infection. Exogenous expression of miR-504-3p, miR-574-5p and miR-1236-5p enhanced HCV-RNA replication 50-, 20- and 5-fold, respectively, while that of mutants possessing a single mutation in the GAGUG motif, miR-504-3p-mt6, miR-574-5p-mt5 and miR-1236-5p-mt5 (Fig 2B), exhibited no enhancement (Fig 2C). Compared with miR-122, the candidate miRNAs have a mismatch sequence at nucleotide positions 8 or 1 in the interacting region with HCV-RNA (Fig 2B). Interestingly, mutant miRNAs containing a modified seed sequence to complement that of HCV-RNA, exhibited significant enhancement of HCV-RNA replication; in particular, the miR-574-5p mutant showed enhancement comparable with miR-122 (Fig 2D). These results imply that the candidate miRNAs participate in the enhancement of HCV-RNA replication via their GAGUG-containing region.

**Fig 2.**
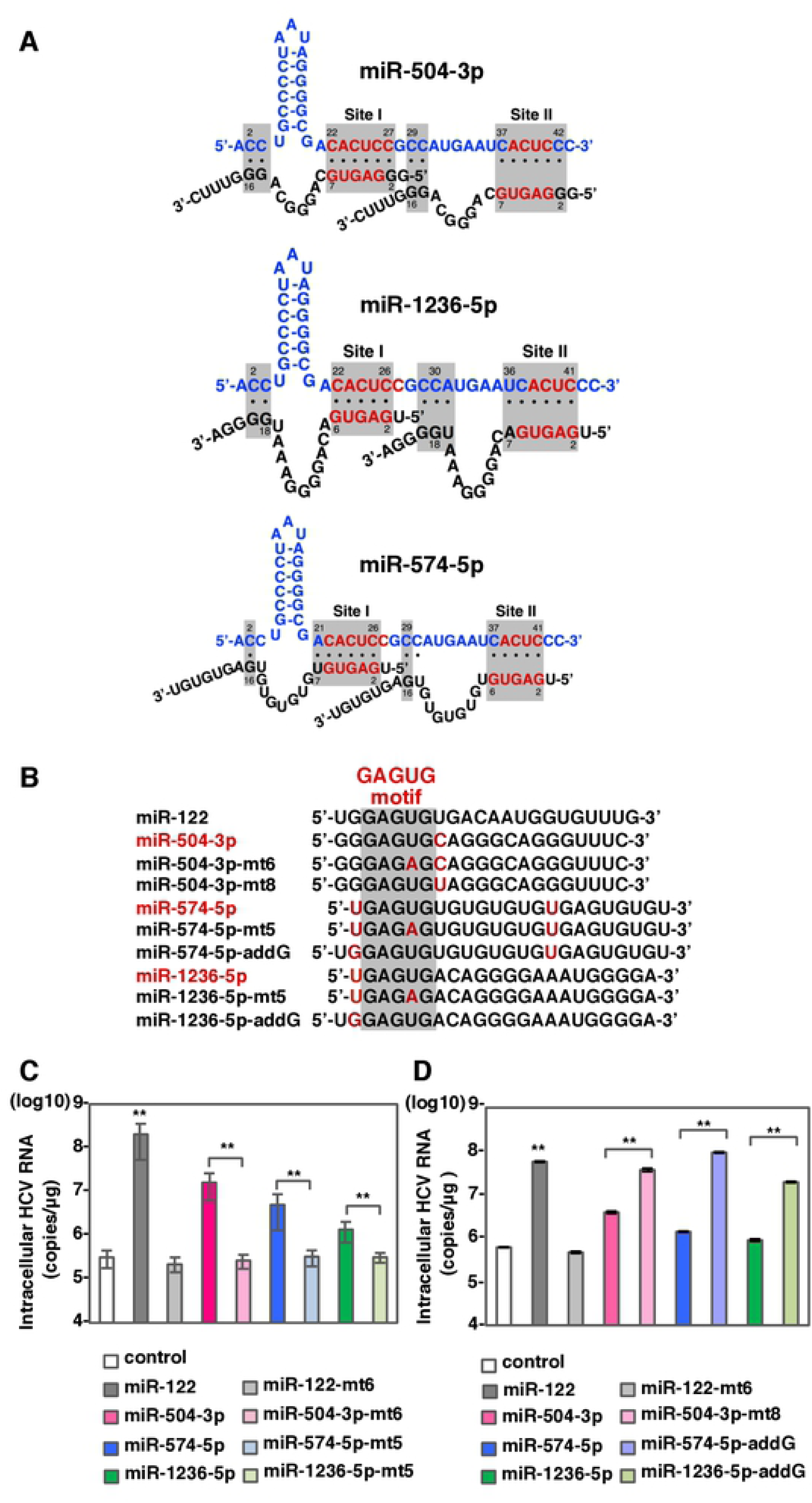
miR-504-3p, miR-574-5p and miR-1236-5p can enhance HCV-RNA replication via miR-122-type interaction. (A) Diagrams of possible interaction between HCV 5’UTR and miR-504-3p, miR-574-5p and miR-1236-5p. (B) Sequence alignment of miR-122, miR-504-3p, miR-504-3p-mt6, miR-504-3p-mt8, miR-574-5p miR-574-5p-mt5, miR-574-5p-addG, miR-1236-5p, miR-1236-5p-mt5 and miR-1236-5p-addG. Mismatching nucleotides were shown in red. (C, D) Intracellular HCV-RNA levels of 751-122KO cells infected with JFH1 in the presence of mimic control, miR-122, miR-504-3p, miR-574-5p and miR-1236-5p or their mutant derivatives were determined at 72 hpi by qRT-PCR. The data are representative of three independent experiments. Error bars indicate the standard deviation of the mean and asterisks indicate significant differences (**P < 0.01) versus the results for the control.

### Engineered non-miR-122-like miRNAs binding to a single site in the 5’UTR are able to enhance HCV-RNA replication

Recently, Schult and colleagues have shown that the interaction of miR-122 with the 5’UTR of HCV alters the structure of the IRES, resulting in the folding of a functional form to promote translation [20]. Based on this report and our observation that candidate miRNAs possessing only a 6-nucleotide match also enhanced the replication of HCV, we hypothesized that modes of targeting HCV 5’UTR positions 23-40 other than that used by miR-122 might exist. Specifically, we examined the possibility of a single miRNA binding site that bridged sites I and II. To test this hypothesis, we prepared mimic miRNAs where a single complimentary site in HCV-RNA would be targeted by 6, 7 or 8 complimentary nucleotides in the miRNA (S4A Fig). 751-122KO cells transfected with each mimic miRNA were infected with HCV and intracellular HCV-RNA was determined at 72 hours post-infection. Among the 6-nucleotide matching mimics we examined, four exhibited slight enhancement of HCV-RNA replication (S4B Fig). In addition, four of the mimics possessing a 7-nucleotide match to HCV-RNA (Fig 3A) enhanced viral replication upon infection with HCV (Fig 3B). Interestingly, not only the mimic possessing a 7-nucleotide match targeting positions 30-36 but also 8-nucleotide match targeting positions 29-36 (Fig 3A) exhibited no effect on HCV-RNA replication upon infection with HCV (Fig 3B). These results suggest that miRNAs binding to a single site masking both G28 and C29 in the 5’UTR with a 7-nucleotide match are able to enhance HCV-RNA replication. We denote this binding mode as “non-miR-122 like”.

**Fig 3.**
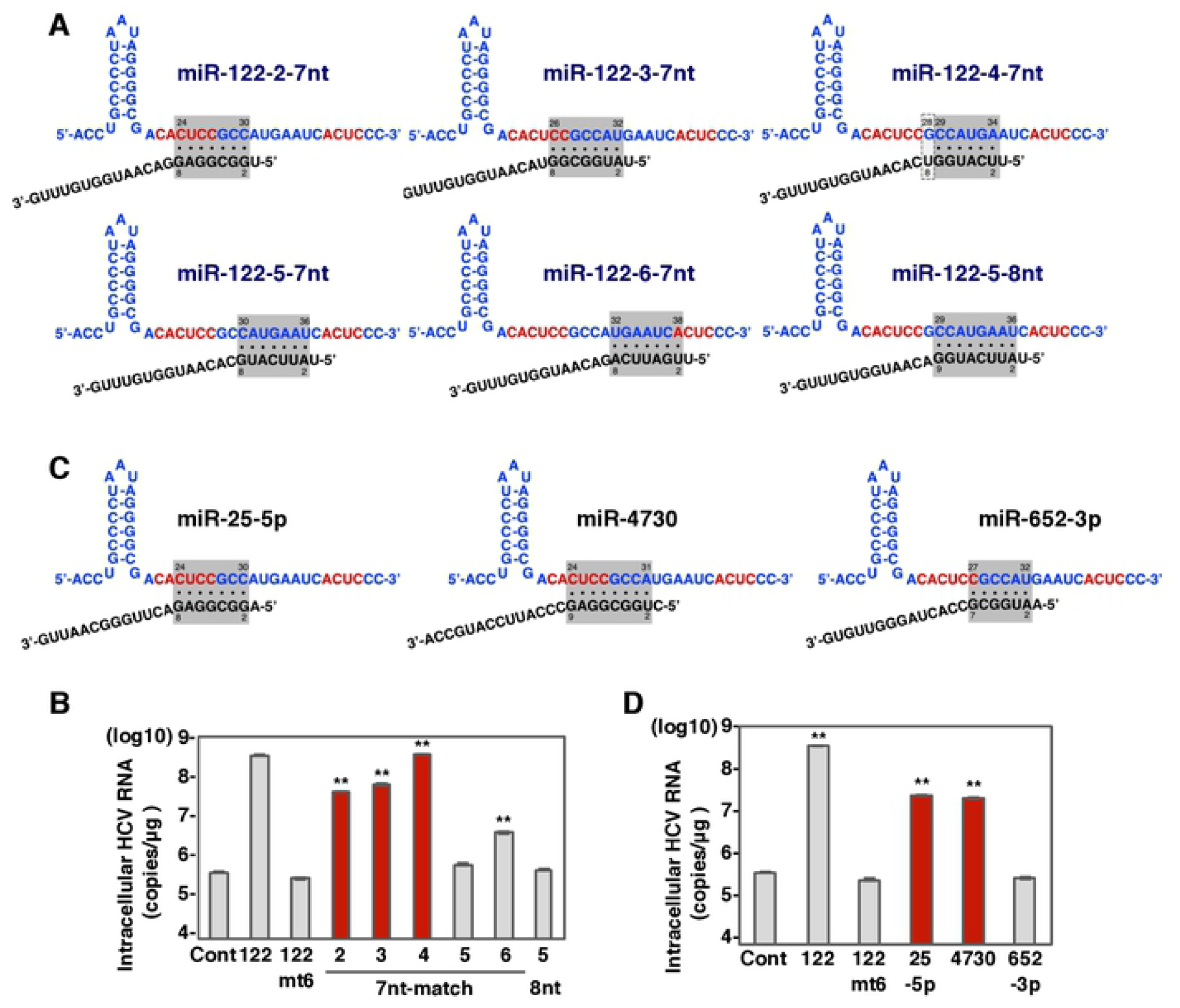
miR-25-5p and miR-4730 can enhance HCV-RNA replication via non-miR-122-type interaction. (A) Diagrams of possible interaction between HCV 5’UTR and synthetic miRNAs with 7nt-match and 8nt-match. Gray-shaded area and framed by dashed line in gray indicate possible interaction region and G-U wobble pair, respectively. Nucleotides in red are important region for the enhancement by miR-122 shown in Fig 1D. (B) Sequence alignment of miR-122 and synthetic miRNAs with 7nt-match and 8nt-match. Mismatching nucleotides were shown in red. (C) Intracellular HCV-RNA levels of 751-122KO cells infected with JFH1 in the presence of mimic control, miR-122, miR-122-mt6 and synthetic miRNAs with 7nt-match shown in Fig 3A. (D) Intracellular HCV-RNA levels of 751-122KO cells infected with JFH1 in the presence of mimic control, miR-122, miR-122-mt6, miR-25-5p, miR-652-5p, miR-1236-5p and miR-4730 were determined at 72 hpi by qRT-PCR. The data are representative of three independent experiments. Error bars indicate the standard deviation of the mean and asterisks indicate significant differences (**P < 0.01) versus the results for the control.

### Natural non-miR-122-like miRNAs enhance HCV-RNA replication

To find natural non-miR-122-like miRNAs, we screened human miRNAs masking G28 and C29 between stem loop I and II of HCV RNA with at least a 7-nucleotide match in the seed region. In this way, two non-miR-122-like candidates (miR-25-5p and miR-4730), with 7 and an 8-nucleotide matches, respectively, were identified (Fig 3C and S3A Fig). In addition, we used miR-652-3p, with a 6-nucleotide match, as a negative control (Fig 3C and S3A Fig). To confirm the activities of each miRNA to suppress translation of target genes, pmirGLO vectors carrying the complementary sequence of each miRNA under the luciferase sequence were transfected into 751-122KO cells. Suppression of luciferase activity was observed in 751-122KO cells transfected with each of miRNA mimics compared to control mimic (S3B Fig). Next, to determine the effect of these miRNAs on HCV-RNA replication, 751-122KO cells transfected with each of the miRNA mimics were infected with HCV and intracellular viral RNA was determined at 72 hours post-infection. Transduction of miR-25-5p and miR-4730 exhibited approximately 60-fold enhancement of HCV-RNA replication, in contrast to that of miR-652-3p, which showed no effect on viral replication (Fig 3D). These results suggest that non-miR-122-like miRNAs binding to single site on HCV-RNA between stem loop I and II via a 7- or 8-nucleotide match have the ability to enhance HCV-RNA replication.

### Expression of miR-122-like and non-miR-122-like miRNAs enhance translation of HCV RNA

Next, to determine the effect of the miRNAs on the translation of HCV RNA, SGR-GND-JFH1-NlucSec RNA, a subgenomic HCV RNA possessing Nanoluc luciferase (NlucSec) gene and a polymerase-dead mutation (S5A Fig), was electroporated into 751-122KO cells together with miRNA mimics. NlucSec activity in 751-122KO cells transduced with mimics of miR-122, miR-122-like (miR-504-3p, miR-1236-5p, miR-574-5p) and non-miR-122-like (miR-25-5p), was higher (in this order), than that of a control mimic (S5B Fig). These results suggest that these miRNAs can facilitate efficient translation of HCV genome.

### Expression of miRNAs other than miR-122 exhibit marginal enhancement on the replication of genotype 1b HCV

Because sequences between stem loop I and II of HCV are different among HCV genotypes and genotype 1b has G28A, G33A and A34G in the stem loop region, we examined the sequence specificity of the enhancement of HCV-RNA replication by miRNAs. miR-6880-5p was identified as a genotype 1b specific non-miR-122 type miRNA (S6A Fig). To examine the enhancement of replication of genotype 1b HCV by miRNAs, 751-122KO cells transfected with mimics of miRNAs (S6A and S6B Figs) were infected with chimeric HCV of genotype 1b Con1 strain and genotype 2a JFH1 strain (Con1C3/JFH) [23]. Among the miRNAs we examined, transfection of mimics of miR-504-3p and miR-574-5p exhibited slight but significant enhancement of replication of the chimeric HCV compared to control mimics, in contrast to large enhancement by that of miR-122 (S6C Fig). These results suggest that expression of miRNAs other than miR-122 exhibit marginal enhancement on the replication of genotype1b HCV.

### Identification of miRNAs that suppress emergence of the G28A mutant of HCV in miR-122-deficient conditions

To examine the emergence of the G28A mutation of HCV during serial passages in the presence of candidate miRNAs, 751-122KO cells transfected with either miRNA mimics of miR-122, miR-122-like (miR-504-3p, miR-574-5p, miR-1236-5p) or non-miR-122-like (miR-25-5p) were infected with HCV and infectious titers in the culture supernatants were determined at each passage (Fig 4A). At passages 3 and 6, we determined the sequence of the 5’UTR of HCV by direct sequencing. The G28A mutation emerged by 6 passages of HCV in 751-122KO cells expressing either control or miR-574-5p mimic, but not in those of other miRNAs (Fig 4B). In addition, expression of miR-122 and miR-574-5p enhanced replication of the G28A mutant of HCV approximately 30- and 4-fold, respectively, probably due to higher affinity than wild type HCV, but that of miR-25-5p, miR-504-3p and miR-1236-5p exhibited no enhancing effect (S7A and S7B Figs). These results suggest that presence of miR-122, miR-25-5p, miR-504-3p and miR-1236-5p suppresses the emergence of the G28A mutation in HCV and that HCV facilitates replication in miR-122-deficient conditions through not only emergence of mutants but also recruitment of miRNAs other than miR-122.

**Fig 4.**
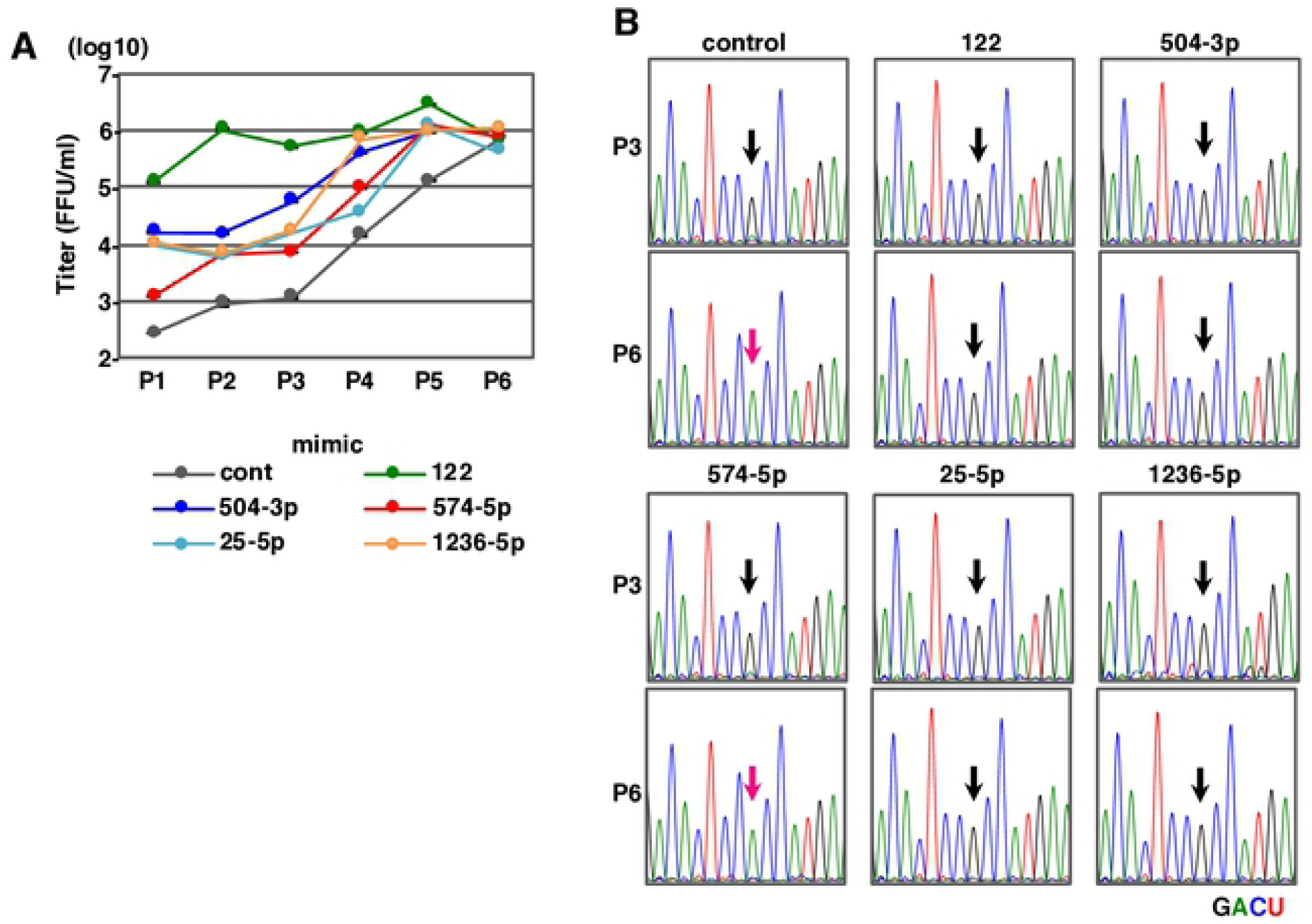
The effect of miRNAs on HCV propagation and the introduction of G28A mutation. (A) Infectious titer in the culture medium on serial passage of 751-122KO cells in the presence of mimic control, miR-122, miR-25-5p, miR-504-3p, miR-574-5p and miR-1236-5p. (B) Mutation of G28A in the 5’UTR of HCV was identified in all independently isolated HCV propagated in Fig 4A. Arrows indicate the position of nt28 in the 5’UTR of HCV. Each RNA base is represented as a colored peak: A, green; U, red; G, black; and C, blue.

### 3D modeling of the miRNA: HCV-RNA:Ago2 complex

In this study, we have shown that not only miR-122-like miRNAs but also non-miR-122-like miRNAs can enhance HCV-RNA translation (S5B Fig) and replication. (Fig 3D). Recent studies have revealed that binding of an miR-122:Ago2 complex or introduction of nucleotide substitutions, such as the G28A mutation, facilitate efficient translation by making an HCV IRES stem-loop (SLII, “long arm”) energetically favorable [20, 21]. More recently, HCV IRES has been reported to hijack the translating 80S ribosome by interaction between the HCV IRES body to the 40S subunit [22]. We hypothesized that the binding of one non-miR-122-like miRNA with a 7-nucleotide match (miR-25-5p) can similarly alter the IRES structure. In contrast, binding of a negative control (miR-652-3p), with only a 6 nucleotide match, is predicted not to alter the IRES structure favorably (Fig 3D). In order to test these hypotheses, we docked representative miRNAs to HCV-RNA, and carried out Replica-Exchange Molecular Dynamics, which can thoroughly sample the resulting conformational space. Specifically, we examined whether miRNA binding would affect the stability of the HCV RNA body or short arm, which are expected to affect HCV RNA binding to the ribosome. 5 representative HCV IRES-miRNA complex models of each representative miRNA molecule were superimposed onto a Cryo-EM reference structure of the HCV IRES bound to the human ribosome, Protein Data Bank (PDB) entry 5a2q, via long arm SLII 39-117 (Fig 5A, top), and the spatial distribution of the body and short arm region were checked by computing the root-mean-square deviation (RMSD) among the models. As shown in Fig 5B, the RMSD of the HCV IRES body and short arm from miR-122 and miR-25-5p were significantly smaller than that of miR-652-3p. Statistical analysis indicated that miR-122 and miR-25-5p were significantly similar, while miR-652-3p was significantly different. This, in turn, suggests that the HCV IRES body and short arm of the miR-652-3p-bound RNA are structurally unstable. In spite of the fact that miR-122 and miR-25-5p interact with HCV-RNA differently, the effect on the stability of the HCV IRES is comparable and distinct from the negative control (miR-652-3p), which interacts with only a 6-nucleotide match. These results indicate that the conformational stability of IRES structure is important for the translational activity of HCV IRES.

**Fig 5.**
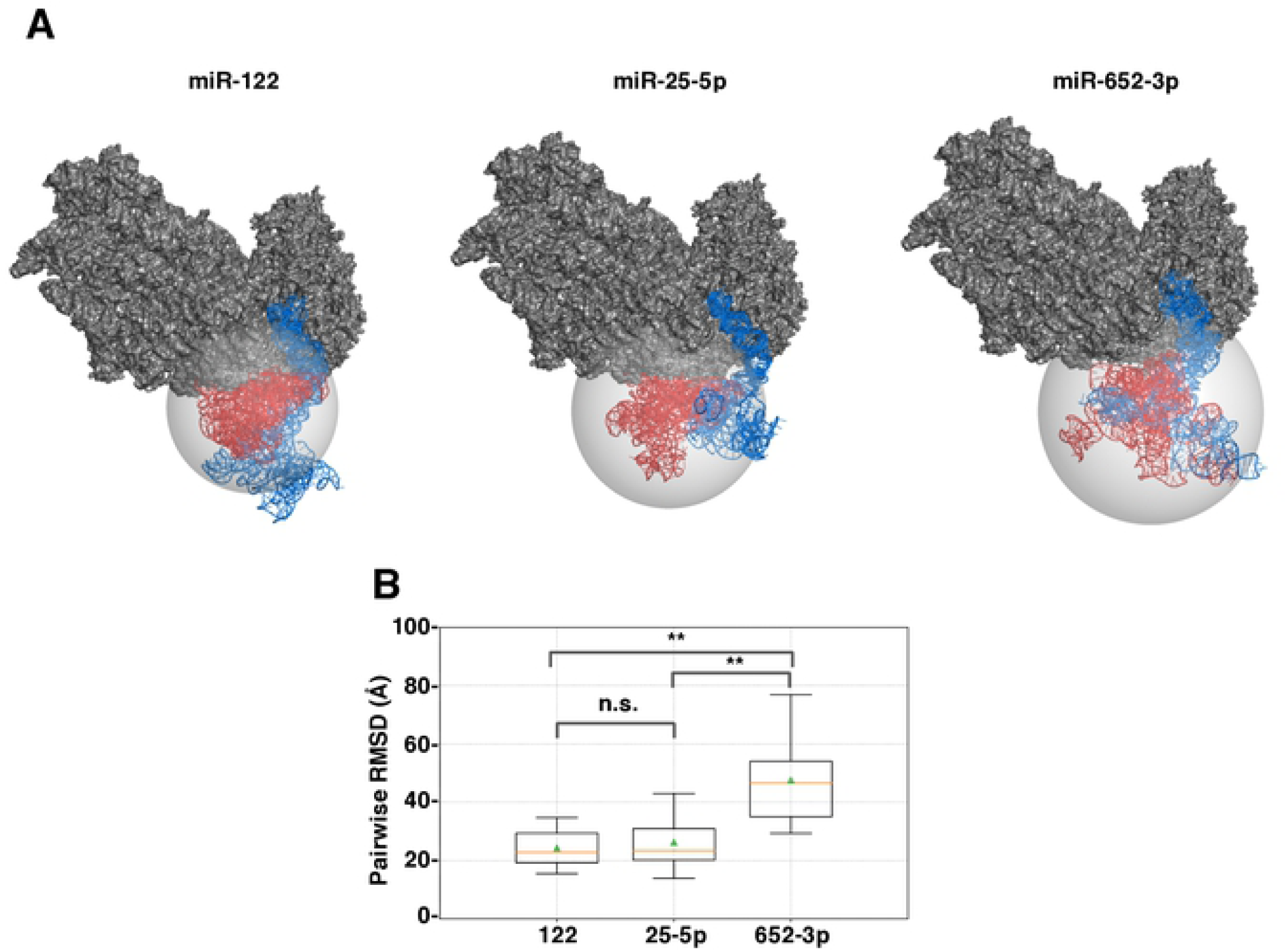
Model of miRNA:HCV-RNA interaction. (A) Superposition results of HCV-RNA tertiary structure onto a Cryo-EM reference structure of the HCV IRES bound to the human ribosome. Representative models of each miRNA (top) were displayed. miRNA binding regions and long arm composed of SLII (nt1-117: blue) and SLIIIa-IIId (nt118-333: red) of HCV-RNA, miRNAs (blue) and 40S rRNA subunit (gray) were shown. Representative models of each miRNA:HCV-RNA:Ago2 (bottom). The distribution of tertiary structures of SLIIIa-IIId were shown as gray sphere. (B) Box-plots of pairwise RMSDs among models of HCV region (SLIIIa-IIId: nt118-333, red blocks in Fig 5A, top). All pairwise RMSDs among 5 representative models (Fig 5A) of nt118-333 regions were determined. The bottom and top boundaries of the box correspond to Q1 (25^th^ percentile) and Q3 (75^th^ Percentile) quartiles, respectively. The lower and upper whiskers correspond to Q1 – 1.5 * IQR and Q3 + 1.5 * IQR, respectively, where IQR = Q3 – Q1. Median value (Q2, 50^th^ percentile) is indicated by an orange line within the box, and mean value is indicated by a triangle symbol in green. Asterisks indicate significant differences (**P < 0.01, n.s.: not significant).

Next, to infer the binding mode of Ago2 on the HCV IRES-miRNA complex model, a complex model was built using PDB entry 6n4o. Specifically, the Ago2 structure was incorporated into the HCV-miRNA models superimposing the double stranded RNA region at the 5’ end of miR-122 and complimentary strand in the HCV-RNA. S8 Fig show alternative binding conformations of Ago2 on different representative HCV-miRNA models. We found that most molecular dynamics snapshots could accommodate Ago2 binding, which is consistent with the previous reports that the formation of SLII of functional IRES is mediated by the interaction with Ago2-miRNA.

### miRNAs other than miR-122 may facilitate HCV-RNA replication in various tissues

Finally, to examine the possibility of participation of miRNAs other than miR-122 in the replication of HCV in non-hepatic cells, expression of the candidate miRNAs in normal tissues was determined by qRT-PCR. Although expression of miR-25-5p, miR-504-3p, miR-574-5p and miR-1236-5p were significantly lower than that of miR-122 in liver, these miRNAs were detected in various normal tissues including brain, thyroid, lung, stomach, small intestine, colon, kidney, liver and bone marrow (S9 Fig). Interestingly, miR-504-3p is expressed mainly in non-hepatic tissues. These observations suggest that not only liver-specific miR-122 but also other miRNAs, such as miR-25-5p, miR-504-3p, miR-574-5p and miR-1236-5p, expressed in various tissues, can facilitate HCV-RNA replication in non-hepatic tissues.

## Discussion

In this study, we examined the possibility of involvement of miRNAs other than liver-specific miR-122 in the enhancement of HCV-RNA replication and showed that HCV-RNA replication is enhanced by the binding of non-hepatic miRNAs. We found that HCV-RNA can be targeted by miR-122-like miRNAs with 6 matching nucleotides containing a GAGUG motif at two sites (sites I and II). Such miRNAs include miR-504-3p, miR-574-5p, miR-1236-5p. In addition, HCV-RNA can also be targeted by non-miR-122-like miRNAs with at least 7 matching nucleotides that can mask G28 and C29 and bind a single site between sites I and II. Such miRNAs include miR-25-5p and miR-4730. Recently, it has been reported that the interaction of miR-122 with the 5’UTR of HCV alters the structure of the IRES, resulting in folding of a functional form to promote translation [20]. Based on the secondary structure of HCV-RNA predicted by selective 2’-hydroxyl acylation analyzed by primer extension (SHAPE), a stem including positions 21-29 was shown to be the minimum free energy structure [20, 35]. In addition, another stem structure including positions 27-33 was predicted as the lowest free energy structure by *in silico* RNA structure prediction algorithms [21]. Although both structures lack stem-loop II (SLII) and fail to form a functional HCV IRES, only a single (G28A) substitution in the 5’UTR of the HCV genotype 2a has been implicated in the folding of SLII without miR-122 binding [20]. These observations suggest that the interruption of base pair of G28 and C29 promotes translation through the formation of SLII. Therefore, it is reasonable to hypothesize that the binding of miRNAs to HCV RNA perturbs the minimum free energy structure, leading to efficient formation of SLII, which, in turn, can be facilitated by the disruption of stem structures including G28 and C29 in the 5’UTR.

Although non-miR-122-like miR-652-3p can interact with the HCV 5’UTR via G28 and C29, which promotes the formation of the SLII structure based on 2D predictions, it did not enhance HCV RNA replication (Fig 3D). We therefore focused on the 3D structural dynamics of the HCV-RNA:miRNA:Ago2 complex. Our 3D structural modeling suggested the importance of conformational stability of the IRES body structure for efficient translation (Fig 5A and 5B). The identification of the HCV IRES tertiary structure by Cryo-EM revealed that it consists of three parts: a long arm (SLII), a body and a short arm [22]. The interaction of the body with the 40S subunit and the structure of the long arm are important for the initiation of translation of HCV IRES [22]. It has been shown that HCV IRES captures the actively translating 80S ribosome and remains bound to the 40S subunit via interactions with the body after translation termination of the 5’-capped mRNA; HCV IRES subsequently places its downstream RNA onto the 40S subunit using the long arm [22]. Based on these observations and our modeling results, it appears that not only the formation of SLII in the HCV 5’UTR, but also alteration of structural dynamics, especially the conformational stability of body, as mediated by the binding of miRNA with Ago2, is required for functional IRES formation. Further studies are needed to clarify the stabilizing effects of non-miR-12-like miRNAs on the HCV IRES structure.

We also observed that, in the presence of either miR-122-like or non-miR-122-like miRNAs, the emergence of the G28A mutation in HCV was suppressed (Fig 4). Our previous report showed that wild type G28 virus was still detected in miR-122-deficient PBMCs from patients infected with genotype 2a HCV and that the G28A mutant exhibited lower replication than wild type in the presence of miR-122 [23]. These observations suggest that the G28A mutant interacting with an Ago2-miRNA complex, mediated by not only miR-122 but also non-hepatic miRNAs, has disadvantages in stages of the viral life cycle other than translation, such as replication, assembly or particle production in miRNA-abundant conditions. Although the involvement of miRNAs in viral particle formation has not been well-studied, it has been reported that the binding of miRNAs such as miR-155 and miR-92a to the HIV-1 genomic RNA can increase efficiency of the packaging of the miRNAs into virions [36]. Because a high level of miRNA incorporation significantly inhibited HIV-1 replication and virion infectivity, it was hypothesized that RNA viruses such as HIV-1 have evolved to avoid cellular miRNA binding to their genome [36]. Other groups have also reported that the regulation of HIV latency by several miRNAs in CD4^+^ T cells and the their inhibition resulted in the enhancement of viral protein expression and particle production [37]. On the other hand, a cis-acting element for HCV-RNA packaging process has been identified as the 3’X region consisting of stem-loop I, II and III in 3’UTR [38] but not in the 5’UTR [39]. Moreover, G28A mutant can easily form SLII in miRNA-independent manner, which is advantageous for translation step, in contrast, it might be not suitable for the replication or particle formation step in the presence of miR-122. Further studies are needed to determine the role of the interaction of miRNA with HCV RNA and the advantages of guanine at 28 in genotype 2a HCV in miR-122-abundant conditions, especially focusing on post-translation step of HCV life cycle.

From an evolutionary perspective, among seven genotypes of HCV, gt2 is predicted to be the oldest lineage, followed by genotypes of 3, 5, and 6 [40]. On the other hand, genotypes of 1 and 4 emerged more recently. Furthermore, characteristic distribution of HCV genotypes different geographical regions [41]. Accordingly, sequences around the miR-122 interaction sites are different among genotypes of HCV [13], for example, the nucleotide position 28 of HCV genome is guanine and adenine in genotype 2 and genotypes of 1, 3, and 4, respectively. These differences of 5’UTR sequence may affect the dependence on miRNAs for translation. Supporting that, we observed the replication of chimeric HCV of genotypes 1 and 2, Con1/JFH, was also enhanced by miRNAs other than miR-122 in genotype 2a JFH-1 strain, while the enhancing effect on replication of Con1/JFH by miR-122-like miRNAs was significantly lower than that by miR-122 (Fig 2C and S6C Fig). In addition, although mutagenesis analysis of the miR-122 binding site in genotype1a H77 strain has shown that the overhanging nucleotide positions of 15 and 16 in miR-122 are required for efficient replication of HCV [14], in this study, we demonstrated that interaction with nucleotide positions 15 or 16 in miR-122 are dispensable for the enhancement of replication of genotype 2a JFH-1 strain (Fig 1C). These observations suggest that the dependence on miRNAs for the enhancement of HCV-RNA replication is different among genotypes, which may be a survival strategy acquired during HCV evolution.

Furthermore, enhancement of HCV-RNA replication by miRNAs other than miR-122 might be involved in the genotype-specific propagation and pathogenesis by the “sponge effect” [19], which is caused by the sequestration of miRNAs by the replication of viral genome, leading to the de-repression of the host target mRNAs. Amyloid precursor protein regulates neurogenesis by antagonizing miR-574-5p in the developing cerebral cortex which promotes neurogenesis [33]. miR-574-5p also participates in the promotion of metastasis and invasion of lung cancer through PTPRU (protein tyrosine phosphatase, receptor type U) [42, 43] and TLR9 signaling [44]. In addition, miR-25-5p has been shown to inhibit cell proliferation of colorectal cancer through activation of AMP-activated protein kinase (AMPK) signaling by silencing protein kinase C ζ [45], suggesting that sequestering of miR-25-5p activates cancer cell proliferation and affects regulation of energy metabolism and maintenance of glucose homeostasis mediated by AMPK signaling [46]. Further studies are needed to clarify the roles of sequestration of miRNAs such as miR-574-5p and miR-25-5p by the replication of HCV in the development of extrahepatic manifestations.

In this study, we have shown the possibility of replication of HCV through interaction with miRNAs other than miR-122 in non-hepatic cells, which is required for the conformational change and stabilization of IRES tertiary structure. These observations provide those miRNAs may lead to the development of extrahepatic manifestations in chronic hepatitis patients through persistent infection of HCV. And we also demonstrate the importance of the 3D structure prediction of IRES through the simulation of HCV RNA:miRNA:Ago2 complex. This technique can be a powerful tool for design of novel anti-HCV drug or treatment targeting HCV-specific system to hijack host translational machinery.

## Materials and Methods

### Plasmids

pCSII-EF-WT-miR-122 and pCSII-EF-AcGFP were described previously [23], respectively. Various mutants of pHH-JFH1mt (pHH-JFH1-E2p7NS2mt; described in [23]) were established by the introduction of a point mutation in miR-122 binding sites of 5’UTR. The complementary sequences of miR-122-5p miR-25-5p, miR-504-3p, miR-574-5p, miR-1236-5p, miR-652-5p, miR-4730 and miR-8074 were introduced into the multicloning site of the pmirGLO vector (Promega, Tokyo, Japan). The plasmid pX330 (Addgene plasmid 42230) designed for the CRISPR-Cas9 system [25, 26] was provided by Addgene. pSGR-GND-JFH1 was established by the introduction of a point mutation of D2760N (from GDD to GND) located in the NS5B polymerase coding region of the pSGR-JFH1. The secreted Nanoluc (Nlucsec) fragment from the pNL1.3 vector (Promega, Madison, WI) was replaced with the neomycin gene of pSGR-GND-JFH1 and the resulting plasmid was designated pSGR-GND-JFH1-Nlucsec. The plasmids used in this study were confirmed by sequencing with an ABI PRSM 3130 genetic analyzer (Life Technologies, Tokyo, Japan).

### Cells, mimic miRNAs and transfection

Human hepatocellular carcinoma cell line Huh7, human embryonic kidney cell line 293T was obtained from Japanese Collection of Research Bioresources Cell Bank (JCRB0403 and JCRB9068). The Huh7-derived hepatocellular carcinoma cell line Huh7.5.1 cells were provided by F. Chisari. All cell lines were maintained in Dulbecco’s modified Eagle’s medium (DMEM) (Sigma, St. Louis, MO) supplemented with 100 U/ml penicillin, 100 µg/ml streptomycin and 10% fetal bovine serum (FBS), and cultured at 37°C under the conditions of a humidified atmosphere and 5% CO_2_. Cells were transfected with the plasmids by using *Trans* IT LT-1 transfection reagent (Mirus, Madison, WI) according to the manufacturer’s protocols. All mimic miRNAs were obtained from Gene Design (Osaka, Japan) and transfected into cells using Lipofectamine RNAi MAX (Life Technologies) according to the manufacturer’s protocol.

### Viruses

pHH-JFH1-E2p7NS2mt was transfected into Huh7.5.1 cells, and the culture supernatants were collected after serial passages. Infectivity of HCV was determined by focus-forming assay and expressed in focus-forming units (FFU) [27]. Unless otherwise noted, cells were infected with HCV at a multiplicity of infection of 1. The lentiviral vectors and ViraPower Lentiviral Packaging Mix (Life Technologies, San Diego, CA) were co-transfected into 293T cells and the supernatants recovered at 48 h post-transfection were centrifuged at 1000 x g for 5 min and cleared through a 0.45 µm filter. The infectious titer of lentivirus was determined by a Lenti-X^™^ qRT-PCR Titration Kit (Clontech, Mountain View, CA).

### Total RNAs from normal tissues and quantitative RT-PCR

Total RNAs from normal tissues such as brain, thyroid, lung, stomach, small intestine, colon, kidney, liver and bone marrow were obtained from Biochain (Newark, CA) and those from cell lines were prepared by using an PureLink^®^ RNA mini kit (Thermo Scientific). Quantitative RT-PCR was performed by using TaqMan^®^ RNA-to-Ct™ 1-Step Kit and a ViiA7 system (Thermo Scientific) according to the manufacturer’s protocol. For quantitation of miRNA, total RNA was prepared from cells by using an miRNeasy mini kit (Qiagen) and mature miR-122-5p, miR-504-3p or miR-574-5p were reverse transcribed by using TaqMan™ Advanced miRNA cDNA Synthesis Kit (Thermo Scientific) and then amplified by using specific primers provided in the TaqMan Advanced miRNA Assays (Thermo Scientific) according to the manufacturer’s protocol. miR-25-3p was used as an internal control [28]. Fluorescent signals were analyzed by using a ViiA7 system.

### Analysis of the 5’UTR sequence of HCV

For a rapid identification of 5’UTR sequence of HCV, RNAs extracted from 100 µl of virus-containing supernatants or PBMCs were amplified by using a PrimeScript^®^ RT reagent Kit (Perfect Real Time) (Takara Bio) and 5’RACE was performed by using a 5’RACE System for Rapid Amplification of cDNA Ends, Version 2.0 (Life Technologies) as described by Ono et al. [23].

### *In vitro* transcription and RNA electroporation

The plasmids pSGR-GND-JFH1-Nlucsec and pJFH1-E2p7NS2mt were linearized by XbaI and transcribed *in vitro* by using the MEGAscript T7 kit (Life Technologies) according to the manufacturer’s protocol. Capped and polyadenylated firefly luciferase (Fluc) RNAs were synthesized by using a mMESSAGE mMACHINE T7 Ultra kit (Life Technologies) according to the manufacturer’s protocol. The *in vitro* transcribed RNA from pSGR-JFH1 (5 µg) was electroporated into cells at 5×10^6^ cells/0.4 ml under conditions of 220V and 950 µF using a Gene Pulser^™^ (Bio-Rad, Hercules, CA) and plated on DMEM containing 10% FBS.

### Analysis of HCV IRES translation activity

*In vitro* transcribed JFH1-NlucSec GND-SGR RNA (5 µg) was electroporated together with Fluc RNA (2 µg) as described above and plated on DMEM containing 10% FBS. Each 3 h post-electroporation, the culture media were harvested and Nanoluc activity was determined in 10 µl aliquots of culture medium using a Nano-Glo^™^ Luciferase Assay System (Promega). To normalize the electroporation efficiency, each cell was lysed in 50 µl of reporter lysis buffer (Promega) at 9 h post-electroporation and luciferase activity was measured in 10 µl aliquots of the cell lysates using a Luciferase Reporter Assay System (Promega).

### Luciferase assay

Cells seeded onto 48-well plates at the concentration of 3 × 10^4^ cells/well were transfected with 250 ng of each of the pmirGLO plasmids and were stimulated with the appropriate ligands for 24 h at 24 h post-transfection. Cells were lysed in 100 µl of passive lysis buffer (Promega) and luciferase activity was measured in 3 µl aliquots of the cell lysates using a Dual-Luciferase Reporter Assay System (Promega). Firefly luciferase activity was normalized with that of Renilla luciferase.

### Docking simulation and modeling

First, we established the secondary structures of HCV-RNA (residues 1-289), miRNA-122 (Fig 1), miR-504-3p (Fig 2), miR-1236-5p, miR-574-5p, miR-25-5p (Fig 3), miR-4730 and miR-652-3p, respectively. In order to predict the intact structures of HCV-miRNA complex, we utilize the cryo-EM structure of HCV (PDB ID: 5A2Q) as restraints to perform iFoldRNA simulation [29, 30]. iFoldRNA is an automatic and accurate RNA 3D structure prediction tool, which is developed on the basis of discrete dynamics simulation (DMD) [31, 32]. Distances between C1’ atoms in 5A2Q are extracted from its 3D structure as restraints to facilitate the RNA modeling by iFoldRNA. iFoldRNA first builds a ring structure composed of all residues in the complex. Then, it conducts a 10000-step initial molecular dynamics simulation to build a preliminary structure. Next, iFoldRNA utilizes replica exchange molecular dynamics simulation (REMD) to thoroughly search through the conformation space. The temperatures of the 8 threads dispatched in REMD are distributed from 0.2 to 0.4 kT. Subsequently, the structures extracted from all 8 trajectories generated by REMD are ranked based on their respective free energies, and the 1% lowest free energy structures are clustered with a specific RMSD cutoff, which is typically one-tenth the number of residues. Since iFoldRNA utilizes a coarse-grained model to expedite the simulation process, the centroids of these clusters will be subject to all-atom reconstruction processes to generate the final all-atom candidate structures of the HCV-miRNA complex.

### Statistical analysis

The data for statistical analyses are the average of three independent experiments. Results were expressed as the means ± standard deviation. The significance of differences in the means was determined by Student’s *t*-test.

## Acknowledgements

We thank M. Tomiyama for her secretarial work, M. Ishibashi, K. Takeda for their technical assistance and M. Hijikata, T. Wakita, R. Bartenschlager, F. Chisari, and M. Whitt for providing experimental materials.

